# A twisted visual field map in the primate cortex predicted by topographic continuity

**DOI:** 10.1101/682187

**Authors:** Hsin-Hao Yu, Declan P. Rowley, Nicholas S.C. Price, Marcello G.P. Rosa, Elizabeth Zavitz

## Abstract

Adjacent neurons in visual cortex have overlapping receptive fields within and across area boundaries, an arrangement which is theorized to minimize wiring cost. This constraint is thought to create retinotopic maps of opposing field sign (mirror and non-mirror representations of the visual field) in adjacent visual areas, a concept which has become central in current attempts to subdivide the cortex. We modelled a realistic developmental scenario in which adjacent areas do not mature simultaneously, but need to maintain topographic continuity across their borders. This showed that the same mechanism that is hypothesized to maintain topographic continuity within each area can lead to a more complex type of retinotopic map, consisting of sectors with opposing field sign within a same area. Using fully quantitative electrode array recordings, we then demonstrate that this type of map exists in the primate extrastriate cortex.

---

Sensory cortices represent the world in mosaics of topographically organized maps. In the visual cortex, neurons in adjacent columns have receptive fields that represent overlapping regions of the retina, both within and across area boundaries. This characteristic is believed to derive from a strong developmental constraint for minimizing the wiring cost of the underlying circuits ^1, 2^. In early visual cortex (e.g. the first and second visual areas, V1 and V2) this results in areas being organized as alternating mirror and non-mirror representations of the visual field ^3, 4^. Indeed, “field sign”, a mathematical property that defines whether a map forms a mirror or non-mirror representation of the visual field ^5^, has become extensively used as a criterion to parse areas in functional mapping studies using both functional MRI and electrophysiology ^6-10^. However, optimizing representational adjacency throughout the cortex is a significant developmental challenge, which current models have only started to address. In primates the arrangement of visual areas rapidly deviates from the concentric V1-V2 configuration towards a tiled arrangement, whereby multiple maps adjoin a single, larger map ^4, 8, 9^. Moreover, different areas develop asynchronously ^4, 11^; thus, a given map may need to organize under a constraint represented by a pre-existing or developmentally more advanced topographic organization along its border.

In a traditional view, each visual area is thought to form a complete and systematic map of the visual field ^12^. However, frequent reports of fractured and incomplete maps have come to suggest that this idealized view seldom applies to extrastriate areas ^13^. Is the universal assumption that each visual area forms a map characterized by uniform field sign also justified? Here, we present a neurodevelopmental model of adjacent cortical areas, constrained only by principles of receptive field smoothness within an area and congruence across area boundaries, which predicts the emergence of two surprising features. First, a “twist” in visual field topography, leading to sectors of mirror- and non-mirror representation within a single area; and second, regions of rapid change in receptive field position within the map, which bridge the transition between mirror- and non-mirror sectors. Using unbiased, high-resolution maps of the controversial dorsal extrastriate region ^14, 15^ we then demonstrate that this type of map actually exists in the visual cortex of a non-human primate (the marmoset). This demonstrates that even under simple wiring principles, the resulting topography of cortical maps can be complex, and suggests that robust parcellation of the visual cortex requires a more detailed analysis than that afforded by any single metric such as field sign.

We focused on the extrastriate region of the primate visual cortex, where the concentric arrangement of V1 and V2 gives way to a patchwork of smaller areas ^8, 9, 13, 16-18^. V2 forms a relatively large but simple map of the visual field where the upper half of the visual field (V2+) is represented ventrally in the cortex, the lower half (V2-) dorsally, and neurons along its rostral border represent the horizontal meridian of the visual field ^3^. What would the retinotopy of a smaller visual area adjacent to V2 look like, if it were to develop representations of both quadrants on the brain’s dorsal surface, constrained by minimum-wiring length principles?

To address this question, we simulated the formation of a retinotopic map adjacent to the rostral border of dorsal V2, by extending a general framework for modelling the development of cortical topographic maps (elastic net ^1, 19^). In this model, “extrastriate” neurons were distributed on a 2D grid representing the surface of the cortex, and their receptive field centers were optimized to cover a set of points distributed regularly in the visual field. The optimization was constrained by two terms in the cost function: the β_1_ parameter governing map smoothness (increasing β_1_ prioritizes matching receptive field locations of neighboring neurons), and the β_2_ parameter governing congruence (increasing β_2_ prioritizes matching the receptive field locations between V2- and the area rostral to V2, at the area boundary).

We found that as the balance between within-area smoothness and between-area congruence was manipulated, three types of maps emerged (Fig. 1). At moderate levels of β_1_ and β_2_, the developed retinotopy was similar to canonical maps such as those proposed for V1 and V2: the eccentricity and polar angle maps were continuous, and the entire map had the same field sign (Fig. 1a, b). As the congruency constraint (β_2_) became stronger than the smoothness constraint (β_1_), however, the map became divided into two regions with opposite field signs (Fig. 1e, f). At the macroscopic level the representations of the central upper and lower visual fields appeared disjointed (Fig. 1e), and receptive fields of neurons along a line crossing the map yielded an “S”-shaped trajectory (Fig. 1g). This map can be considered a “twisted” version of the first map (compare Fig. 1h with 1d); the twist rotates the direction of the eccentricity gradient, which reverses the polarity of the field sign, and divides the area into two regions that are not mirror images of each other. Given that occipital areas develop asynchronously according to a caudal to rostral gradient ^4, 11^, the assumption that β_2_ may be stronger than β_1_ at specific stages of map maturation rostral to V2 is biologically plausible. Finally, at high values of β_1_ and β_2_, another type of map with opposing field signs developed. In this map, the twist is so severe that the representation of the fovea in the lower field is displaced from the rest of the lower field representation.

**Fig. 1.**
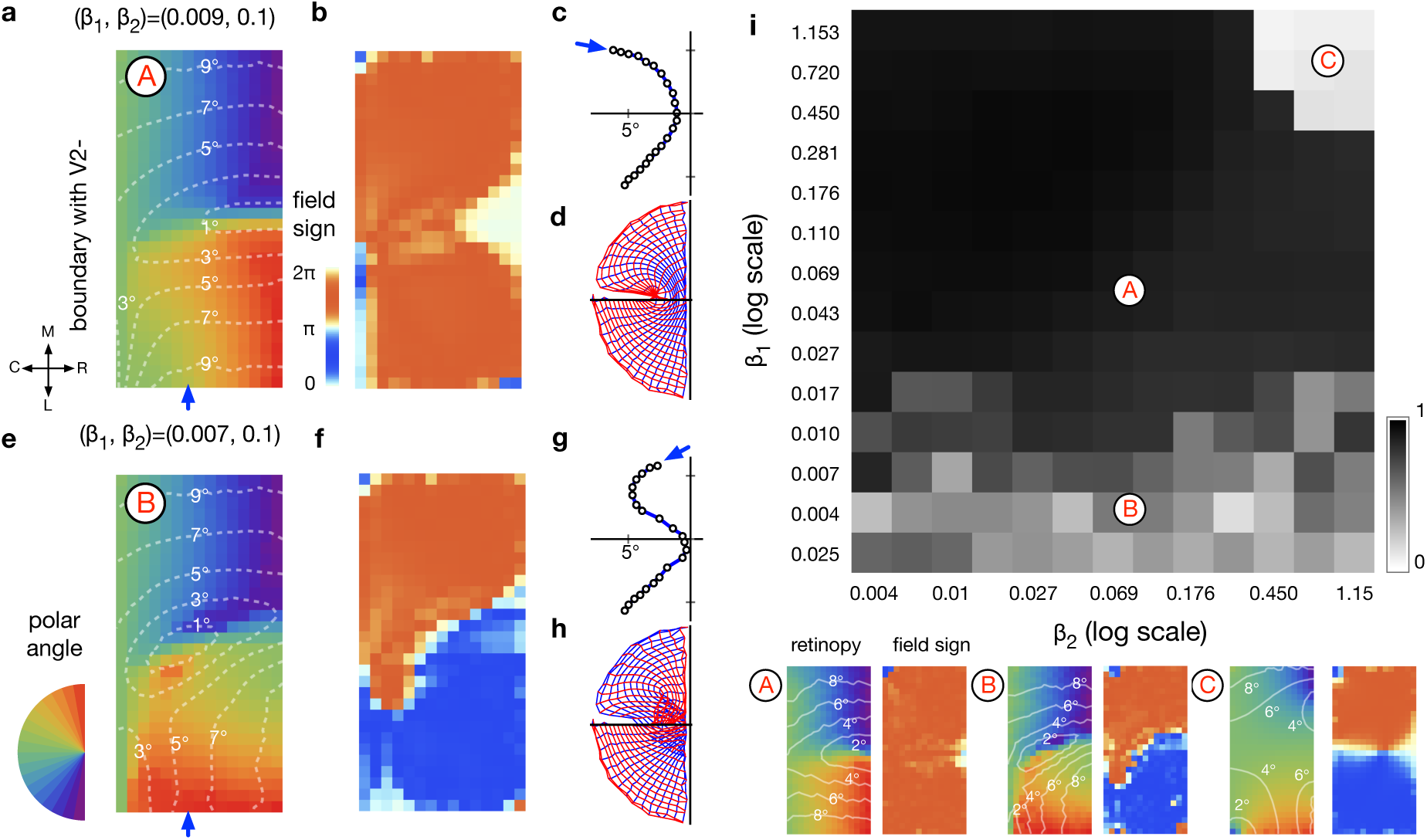
Simulations of extrastriate map formation produce three types of maps depending on how its constraints are prioritized. (a) Moderate levels of β_1_ and β_2_ lead to smooth retinotopy; colors represent polar angle, and dashed white contours represent eccentricity. (b) field sign at each location. (c) Receptive field centers progress in a smooth arc if an “electrode” is moved in the direction indicated by the blue arrow in **a** (d) The grid of the cortical map (blue: columns; red: rows) changes smoothly at all points. (e-h) If smoothness (β_1_) is less strongly weighted, a twisted receptive map emerges, with two field signs and receptive fields that progressed in a “S”-shaped trajectory. The twist in the cortical map grid accounts for the trajectory in c. (i) The dependency between the two parameters and the field sign of the resulting map. The grey scale represents the averaged field sign, scaled to −1 to 1, across the map and across 8 repeats of randomly initialized simulations. It measures the complexity of the maps: values close to 0 indicate maps with balanced regions of opposing field signs, whereas values close to 1 indicate maps dominated by the mirror image field sign. Regions indicated by A, B, and C correspond to the three types of maps illustrated below.

We have shown that it is possible for twisted maps to develop under very basic constraints, but do such maps form in real brains? The retinotopy of the extrastriate region rostral to V2 in non-human primates offers a promising target to test the predictions of the model. Based on histology, connectivity and single-unit electrophysiology, it has been proposed that this region harbors a dorsomedial area (DM) which forms an unusual retinotopic map showing regions of different field sign (Fig. 2a)^17^. However, this proposal was based on qualitative evidence, and the available data also led to other models according to which the same region contains areas with more conventional retinotopy ^8, 14, 15^.

**Fig 2.**
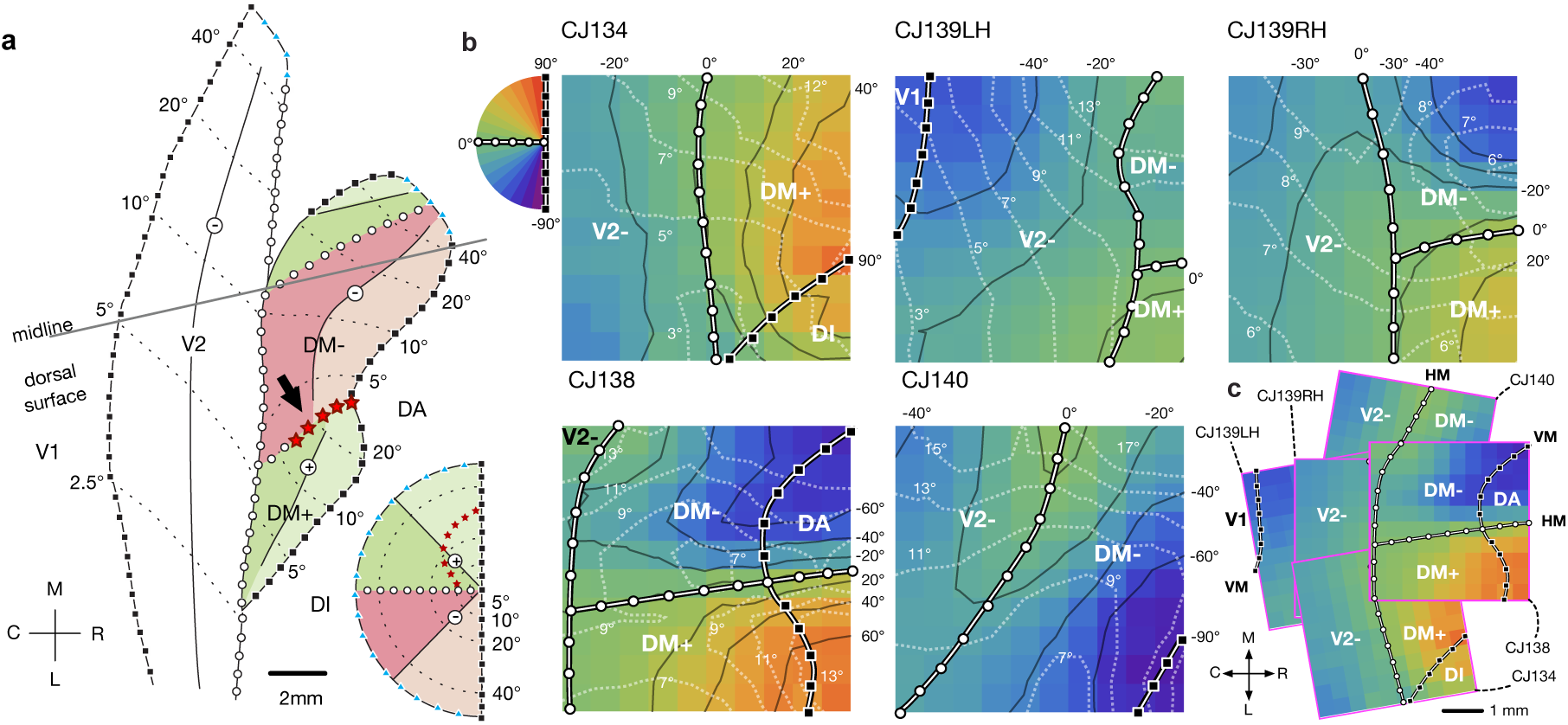
The retinotopy of the dorsomedial cortex. (**a**) A schematic summary of one of the competing models of organization of dorsomedial cortex in the marmoset^9^. The inset at the bottom right illustrates the color scheme used to illustrate different segments of the visual hemifield in the proposed area DM. The arrow indicates the location of the putative “map discontinuity”. (**b)** The retinotopy of 5 hemispheres from 4 animals (identifiers of the cases are prefixed by “CJ”), estimated from receptive fields mapped using 10×10 electrode arrays. The colors represent the polar angles. Polar angles are indicated by solid black contours and numbers in black. Eccentricities are indicated by dashed white lines and numbers in white. Inset: The color scale for representing the polar angle in **b** and **c**. The horizontal meridian (polar angle=0°) is indicated by thick lines overlaid with circles. The vertical meridian (polar angle ±90°) is indicated by thick lines and squares. (**c**) Composite summary of the spatial relationships shown in **b**. Abbreviations: V1: primary visual area; V2-: dorsal portion of secondary visual area; DM+, DM-: upper/ lower field representation of the dorsomedial area; DA - dorsoanterior area; DI – dorsointermediate area; M - medial; L - lateral; C - caudal; R - rostral.

To produce an unbiased, evenly-sampled map, we made quantitative measurements of receptive field locations and response properties of neurons in the region surrounding the rostral border of dorsal V2 (lower quadrant representation, V2-), using 10×10 multielectrode arrays with 400 µm electrode spacing (Fig. 2; Supplementary Figs. S1-5) in 5 hemispheres of 4 marmoset monkeys (*Callithrix jacchus*). Our data indicated that immediately rostral to V2-there is a representation of the contralateral visual field covering at least 20° in eccentricity (Fig. 2b, c), where neurons with receptive fields of similar size (Supplementary Fig. 6a) and response properties (Supplementary Fig. 6b) are found. In agreement with some of the earlier reports in this region the upper field is represented laterally (referred to here as DM+), and the lower field medially (DM-); both DM+ and DM-border V2-along a continuous representation of the horizontal meridian ^16-18^.

Earlier work ^13, 17^ has suggested that the transition between DM+ and DM-includes a “map discontinuity” (arrow in Fig. 2a) – a sudden “jump” of receptive field centers between closely spaced recording sites, which violates topographic continuity. Using quantitative techniques (Fig. 3a), we can see that the representation in this region is topographically unusual, but actually not discontinuous. The position of the receptive field centers changes rapidly (Fig. 3b), following a continuous “S”-shape trajectory across the horizontal meridian (Fig. 3c), similar to the configuration predicted by the modelling (Fig. 1g). However, the receptive fields of adjacent recording sites still overlap substantially, indicating that this is more accurately described as a thin strip of cortex with an eccentricity gradient that rapidly reverses polarity (Fig. 3b). This “S”-shaped trajectory, and the accompanying change in field sign (Fig. 3d, e), demonstrate that the predicted topographical “twist” occurs in nature. However, the data also indicate that this is not achieved at the expense of violating local receptive field adjacency.

**Fig 3.**
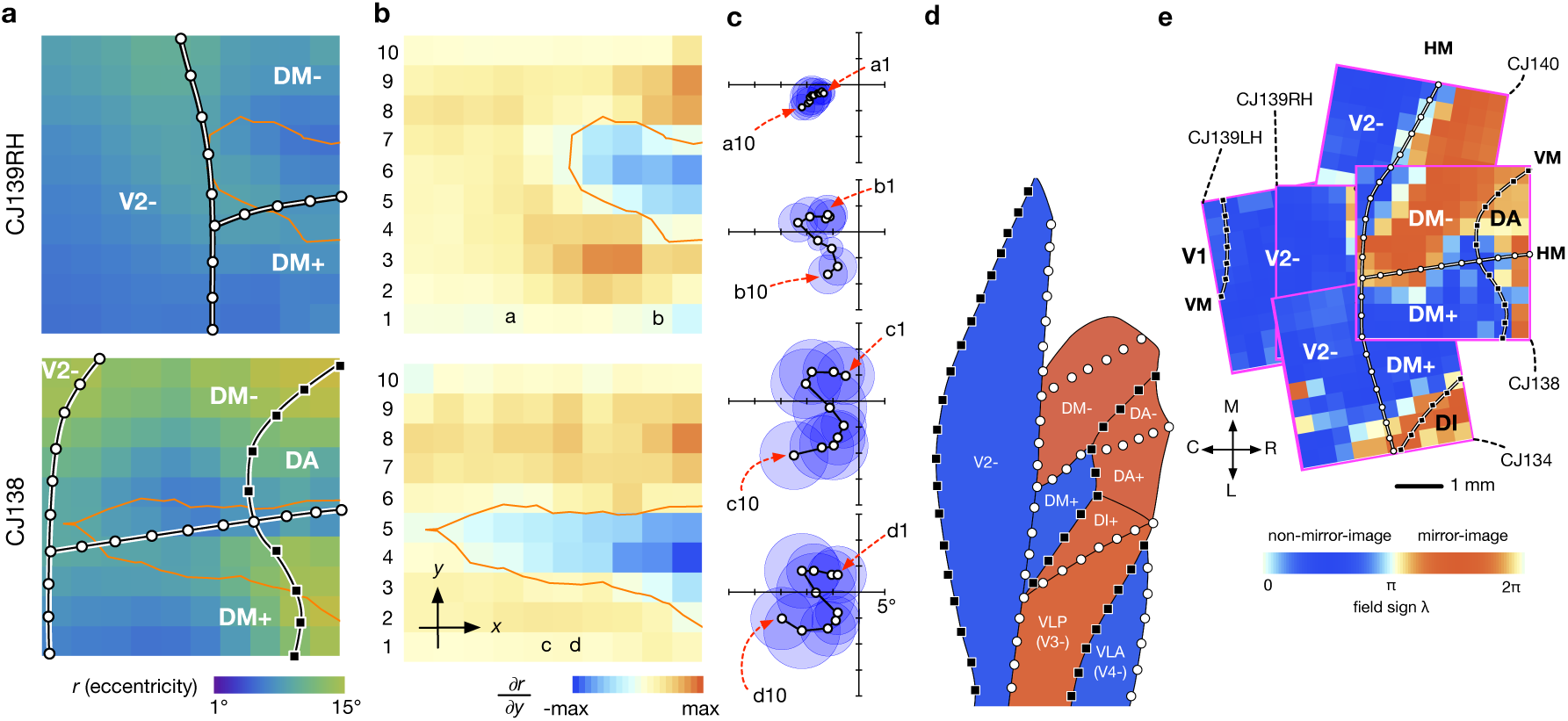
Local features of the DM map. (**a**) Eccentricity maps of two selected cases. The cortex enclosed by the orange contours indicates a region where the gradient of the eccentricity rapidly reversed polarity. (**b**) This region can be visualized by plotting the partial gradient of eccentricity with respect to the spatial dimension of the rows (the y-axis) of the electrode arrays. The region in blue (enclosed by the orange contour) corresponds to sites where the eccentricity of the receptive field rapidly decreased in the lateral-to-medial direction. (**c**) Representative sequences of receptive fields associated with channels in columns of the electrode arrays. The association between the receptive fields and the channels are identified by letters (columns) and numbers (rows). The progressions of receptive fields followed a distinctive “S”-shape trajectory. (**d**) A summary of the field signs for areas in the dorsomedial region of the marmoset visual cortex. The field signs for area DM, DI, VLP (ventrolateral posterior area, or V3) and VLA (ventrolateral anterior area, or V4) were inferred from published maps. **(e)** Field sign maps estimated for the 5 cases.

In summary, we describe an unbiased, finely sampled visual cortex map with a “twist” in its topographic organization, and demonstrate that the unusual retinotopy can arise naturally from the same processes that are thought to explain the organization of early areas such as V1 and V2.

Two types of discontinuity in topographic maps can be distinguished ^13, 20^: *field discontinuities* (i.e., adjacent regions on the visual field mapped to non-adjacent regions on the cortical surface) and *map discontinuities* (i.e., non-adjacent regions on the visual field mapped to adjacent regions on the cortical surface). While field discontinuities are well-documented, and appear ubiquitous in the primate cortex ^3, 7, 13^, the existence of map discontinuities has been controversial. We showed that the topographic transition at the DM+/ DM-boundary is more appropriately characterized as a thin strip of cortex with rapidly changing gradient, rather than a true discontinuity. A map discontinuity has also been reported to exist at the border between hand and face representations in the somatosensory cortex ^21^, but the fine topography of this region is yet to be studied with fully quantitative techniques.

We showed that different types of retinotopic maps can emerge depending on the balance between within-area continuity and between-area congruency. While the developmental factors that determine the strength of these two constraints remain unknown, our simulation indicates that the DM map was formed in a regime where between-area congruency was prioritized over within-area continuity. This situation may emerge when topographic maps in different areas develop asynchronously, with a pre-existing map in an early-maturing area constraining the possible receptive field locations at the border with a late-maturing area ^22^. Congruency allows multiple areas to form large-scale supra-areal clusters, which have been proposed as the fundamental building block of cortical organization ^7, 23^. Studying the consequence of continuity constraints, in the context of the geometrical relationship between areas, is critical for the understanding of these large-scale structures.

Due to the similarity between DM+ and DM-in terms of histology, connectivity, receptive field size, cortical magnification factor, and response properties (Supplementary Fig. S6), the modeling result further supports the notion that these are parts of the same area, which is located immediately rostral to dorsal V2. This organization is unlikely to be unique to the marmoset, as recent studies in owl monkeys ^8^ and macaques ^9^ reported compatible results in the cortex rostral to V2, but subdivided the cortex differently by virtue of prioritizing the unity of field sign within areas. However, both of these proposals resulted in an areas with unbalanced representations of the upper and lower quadrants. Whereas in many situations the areal boundaries can be readily identified by the reversals of field sign, our results show that regions of opposing field sign can arise naturally within a single area based on a similar developmental mechanism.

In summary, the interpretation of field sign for area parcellation demands a nuanced approach, which takes into account the context of the global map across areas, cyto- and myeloarchitecture, and connectivity. The present results indicate that some of the uncertainties and controversies in the mapping of extrastriate areas might be due to complex local topographies occurring at the microscopic level, but being undetected given the typical resolution of fMRI.

## Supporting information

Supplementary Methods and Findings

