## Supplementary Methods and Findings for "A twisted visual field map in the primate cortex predicted by topographic continuity"

**Preparation.** Experiments were conducted in accordance with the Australian Code of Practice for the Care and Use of Animals for Scientific Purposes. All procedures were approved by the Monash University Animal Ethics Experimentation Committee. In four marmoset monkeys (*Callithrix jacchus*) surgical anaesthesia was induced by intramuscular injection of alfaxalone (Alfaxan, 8mg/kg). Under anaesthesia, the animal was injected with an antibiotic (Norocillin, 25mg/kg) and dexamethasone (Dexason, 0.3mg/kg). A tracheotomy was performed, and the femoral artery was cannulated. The animal was positioned in a stereotaxic frame, and a craniotomy and durotomy were performed over the dorsomedial extrastriate cortex. For the duration of the recording session, the animal was maintained on infusion of sufentanil citrate (Sufenta Forte, 250µg/5ml), pancuronium bromide (Pancronium, 4mg/2ml), dexamethasone (Dexapent, 5mg/ml), xylazine (Xylazil-20, 20mg/ml) and salts and nutrients (0.18% NaCl/4% glucose solution, Synthamine-13 and Hartmann's solution), and ventilated with a nitrous oxide and oxygen mixture (7:3). The animal's body temperature was kept at a steady 38 degrees, measured by rectal thermometer. The eye contralateral to the craniotomy was held open, and atropine (Atropt, 1%), phenylephrine, and carmellose sodium (Celluvisc) eye drops applied, before a contact lens was inserted to focus the eye at a viewing distance of 20-40cm. The ipsilateral eye was protected with carmellose sodium, closed, and occluded.

**Electrophysiology.** 10x10 "Utah" arrays (Blackrock Microsystems, Salt Lake City, USA) with 96 active channels, were implanted in the expected location of the border between V2 and the areas rostral to it, using a pneumatic insertion tool. The position of the array was planned using stereotaxic coordinates [24] *in vivo*, and verified with flatmount histology *post mortem* (Supplementary Figure 7). Electrodes were 1.5 mm long, and spaced at 400 µm intervals. The raw voltage signal was recorded at 30kHz using a Cerebus system (Blackrock Microsystems, Salt Lake City, USA) and high-pass filtered at 750Hz. Spikes were detected using automatic thresholding of the local signal. After recording, manual spike sorting was performed offline using Plexon Offline Sorter (Plexon Inc., Dallas, USA).

**Visual stimulation.**

The positions of several receptive fields in space were hand-mapped on a tangent screen, and a VIEWPixx 3D (VPixx Technologies, Saint-Bruno, Canada) positioned at a viewing distance of 350-450 mm with the receptive fields in and around the center of the monitor. The stimuli were presented at a 120Hz refresh rate using The Psychophysics Toolbox in MATLAB [25-27]. Receptive fields were mapped at 1° resolution with both "on" (white) and "off" (black) squares flashed on a grey background. Squares appeared for 100 ms with a 50-to-100 ms (different in different cases) inter-stimulus interval.

**Retinotopy.** To quantify the geometry of the receptive fields, the spike counts elicited by each location of the flashing square stimulus were smoothed with a 5x5 Gaussian kernel. A Gaussian function was fitted to the smoothed map (EQ. 1), where ($\mu_x$, $\mu_y$) is the center of the receptive field. The boundary of the receptive field (and therefore its size) was determined by the contour at 15% of the peak response.

$$r = c + e^{\frac{1}{2}\left(-\frac{(x-\mu_x)^2}{\sigma}-\frac{(y-\mu_y)^2}{\sigma}\right)} \qquad \text{EQ. 1}$$

The center of gaze was inferred from the retinotopy, given that at the boundary of visual areas, the progression of the receptive fields reverses its direction at the horizontal or the vertical meridian. For CJ134, CJ138, CJ140, the locations of the blind spot could be identified in the receptive field maps (Figs. S1, S4, S5). Because the representation of the blind spot on the visual field is approximately 15° away from the fovea on the horizontal meridian [28], this imposed a strong constraint on the location of the center of gaze.

**Field sign.** Field sign (λ) is defined as the clockwise angle between the eccentricity gradient and the polar angle gradient [5]. For calculating the gradients, the coordinates of the receptive field centers were smoothed by moving window averaging. The field signs calculated for individual channels were then smoothed by moving window averaging. For visualization, the calculated field sign was compressed by a sigmoid function[5] and then displayed with a color scale that distinguish non-mirror image maps ($0<\lambda<\pi$) from mirror-image maps ($\pi<\lambda<2\pi$).

**Simulation.** A modified version of the elastic net algorithm [19] was used to solve the proximate minimal path length problem[1]. The model consisted of neurons with point receptive fields ($y_j$) initialized to random locations on the contralateral visual hemifield within 10° of eccentricity. These receptive fields are arranged topologically in a 30x15 grid modeling area DM, and they were updated iteratively using gradient descent to minimize the energy function (EQ. 2):

$$E = -k \sum_{i} log \sum_{j} \Phi\left(x_i, y_j, k\right) + \beta_1 \sum_{j} \sum_{j' \in N(j)} \left\| y_{j'} - y_j \right\|^2 + \beta_2 \sum_{j \in B} \sum_{j' \in N_{V2}(j)} \left\| z_{j'} - y_j \right\|^2 \qquad \text{EQ. 2}$$

The first term is a "coverage term" that forces $y_j$ to converge to 500 fixed points $(x_i)$ distributed regularly on the visual hemifield up to 10° of eccentricity, with a density that dropped off with eccentricity (density ∝ eccentricity$^{-0.4}$). $\Phi(x_i, y_j, k) = exp(\frac{-\left\| x_i - y_j \right\|^2}{2k^2})$, where $k$ is an annealing factor that was initialized to 30.0, and reduced by 0.5% for each iteration. The regular distribution of $x_i$ was implemented with the Vogel method. The coverage term is followed by two regularization terms to enforce topographic continuity, which were weighted by two parameters $\beta_1$ and $\beta_2$. The first regularization term enforces the smoothness of the retinotopy, where $N(j)$ denotes sites on the 30x15 grid that neighbor site $j$. The second regularization term enforces congruency with the retinotopy of V2 at the caudal boundary of DM (denoted by $B$), where $N_{V2}(j)$ denotes sites in V2 which neighbor site $j$ in DM. V2 receptive fields are located at $z_{j'}$, which are fixed points on the horizontal meridian. The range of eccentricity at the DM/V2 boundary was 2° to 10.0°. The model was implemented with Tensorflow and the source code is available at https://github.com/hsinhaoyu/DM_Retinotopy.

**Cortical magnification factor**. The reciprocal of the (linear) cortical magnification factor [29] $1/M$ was calculated as $\sqrt{1/M_a}$, where $1/M_a = |\det(J)|$, $J$ being the Jacobian matrix of the mapping from the cortex surface to the visual field. This measures the linear cortical magnification factor $M$ from the areal cortical magnification factor $M_a$ assuming that the mapping is isotropic. The calculation of $J$ was based on the locations of the measured receptive field centers without smoothing. The estimated $1/M$ was then spatially smoothed.

**Orientation tuning**. We used drifting sinusoidal gratings to determine the preferred orientation for the units on the array. Spatial and temporal frequency were selected to best drive the largest number of units possible and ranged from 0.3 to 1 cycle/° and 2.5 to 4 Hz. Responses were measured for 24 directions, tiling 360° at 15° intervals to motion lasting 400 to 1000 ms. The preferred orientation was determined based on the resultant vector [30], and the bandwidth with the circular variance of the responses [31, 32].

**Histology**. After the completion of data collection, the animal was given a lethal overdose of sodium pentobarbitone (100mg/kg). The array was removed and the animal transferred to a fume hood where it was perfused with buffered saline. The unfixed brain was immediately extracted, and the two hemispheres were separated and physically flat-mounted. Flat-mounting (Fig. S7) was performed by gently dissecting away the white matter of the cortex with dry cotton swabs, with the cortex supported on a piece of moist filter paper (pial surface down). Relaxation cuts were made in the fundus of the calcarine sulcus, and at the anterior end of the sylvian sulcus to allow the cortex to lie flat. The cortex was held in fixative between two large glass slides under a small weight overnight, and then was soaked in sucrose solution in increasing concentrations (10%, 20% and 30%). The flat-mounted hemisphere was then cut in a cryostat to a thickness of 40 μm. Alternate sections were stained for myelin and cytochrome oxidase.

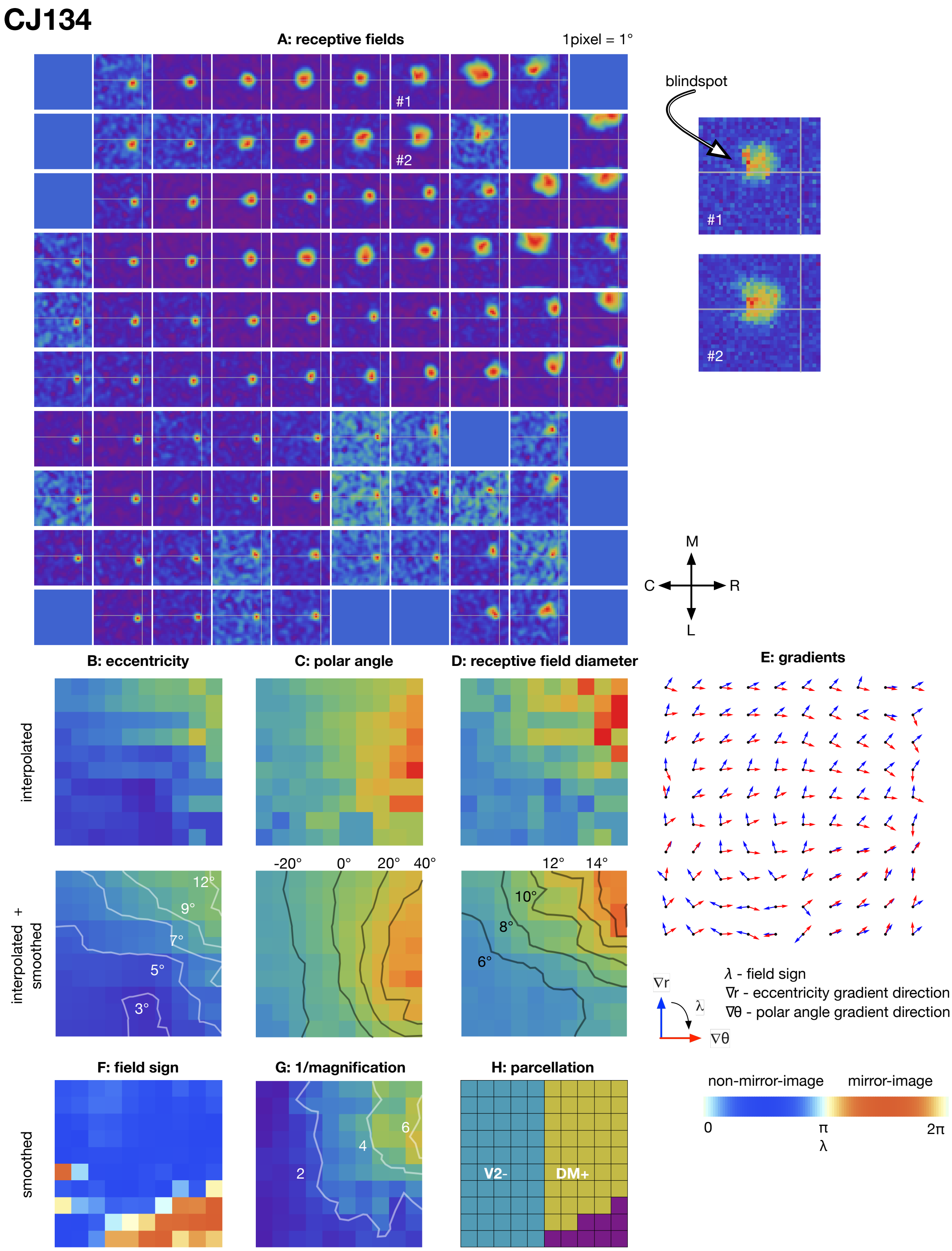


**Supplementary Figure 1**

**Summary mapping data for case CJ134**

**(A)** The outlines of the receptive fields for each channel, measured with a flashing square stimulus. In each plot of the receptive fields, the estimated horizontal and vertical meridians are indicated by thin white lines. For some cases, the representations of the blind spot were visible in some of the receptive field maps. Representative receptive field maps with "holes" (the blind spot) are shown on the right without spatial smoothing. The correspondence between those maps and the plots for the entire array is indicated by numbers in white**. (B, C, D)** Maps of the eccentricity, polar angle, and diameter of the receptive fields without (top) and with (bottom) spatial smoothing**. (E)** The gradient vector fields used to estimate the field sign. The blue vectors represent the directions of the eccentricity gradient ($\nabla r$), and the red vectors represent the directions of the polar angle gradient ($\nabla \theta$). Field sign ($\lambda$) is the clockwise angle between the two vectors. **(F)** The map of field signs, illustrated with the color scale shown in the right**. (G)** The map of the reciprocals of the linear cortical magnification factor (unit: $°/mm$). **(H)** The assignments of the electrode array channels to one of four areas (V2-, DM+, DM-, and others).

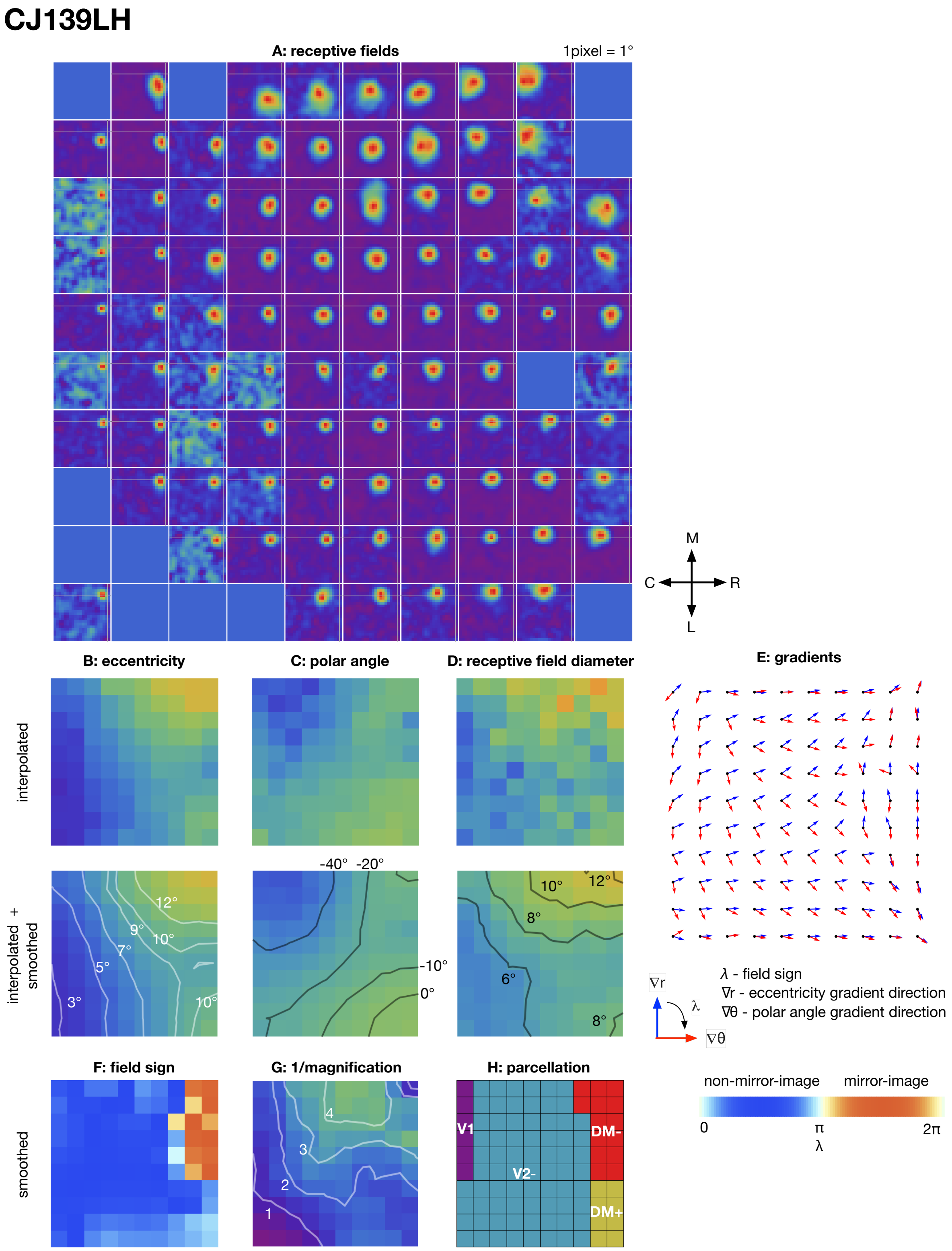


**Supplementary Figure 2**

**Summary mapping data for case CJ139LH**

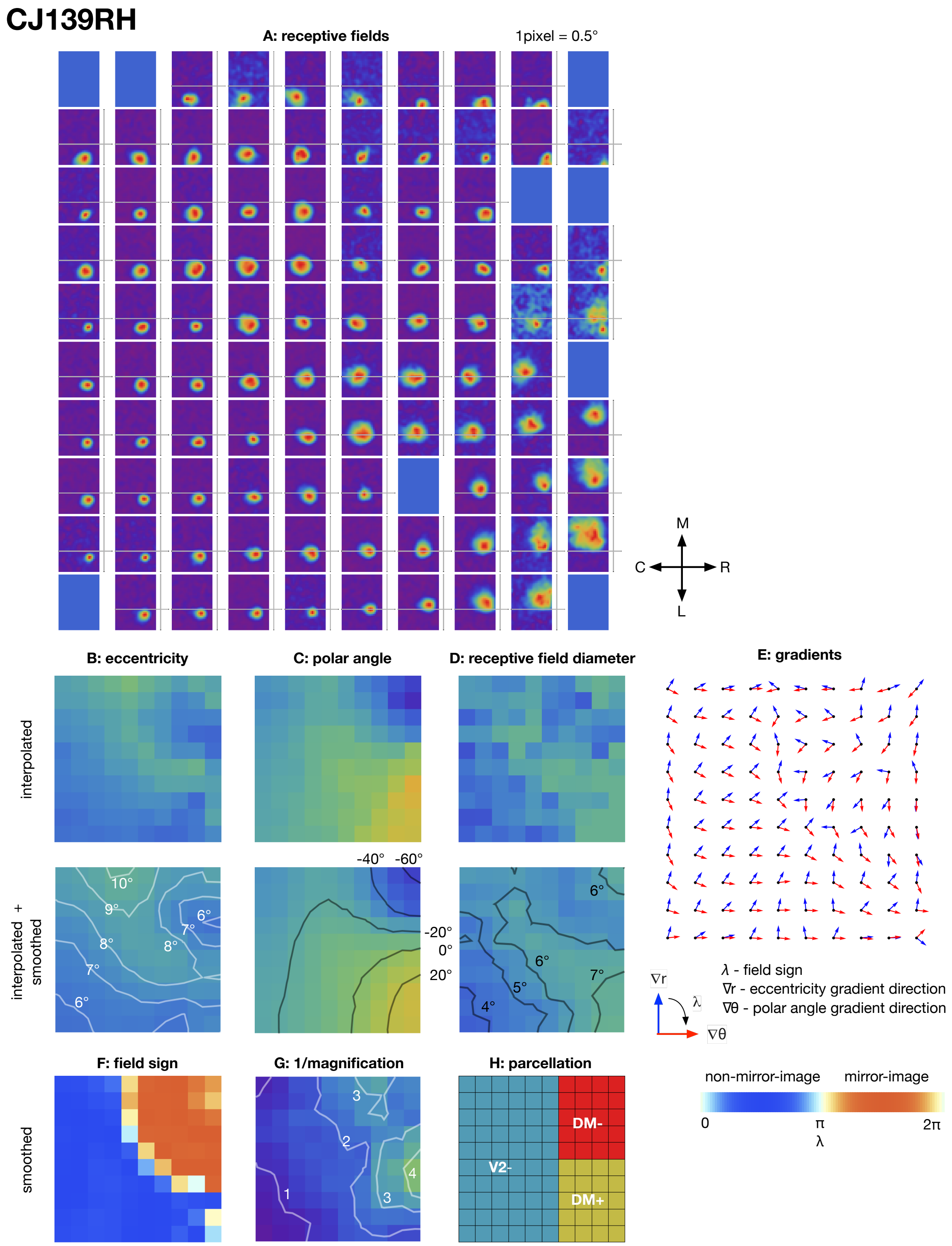


**Supplementary Figure 3**

**Summary mapping data for case CJ139RH**

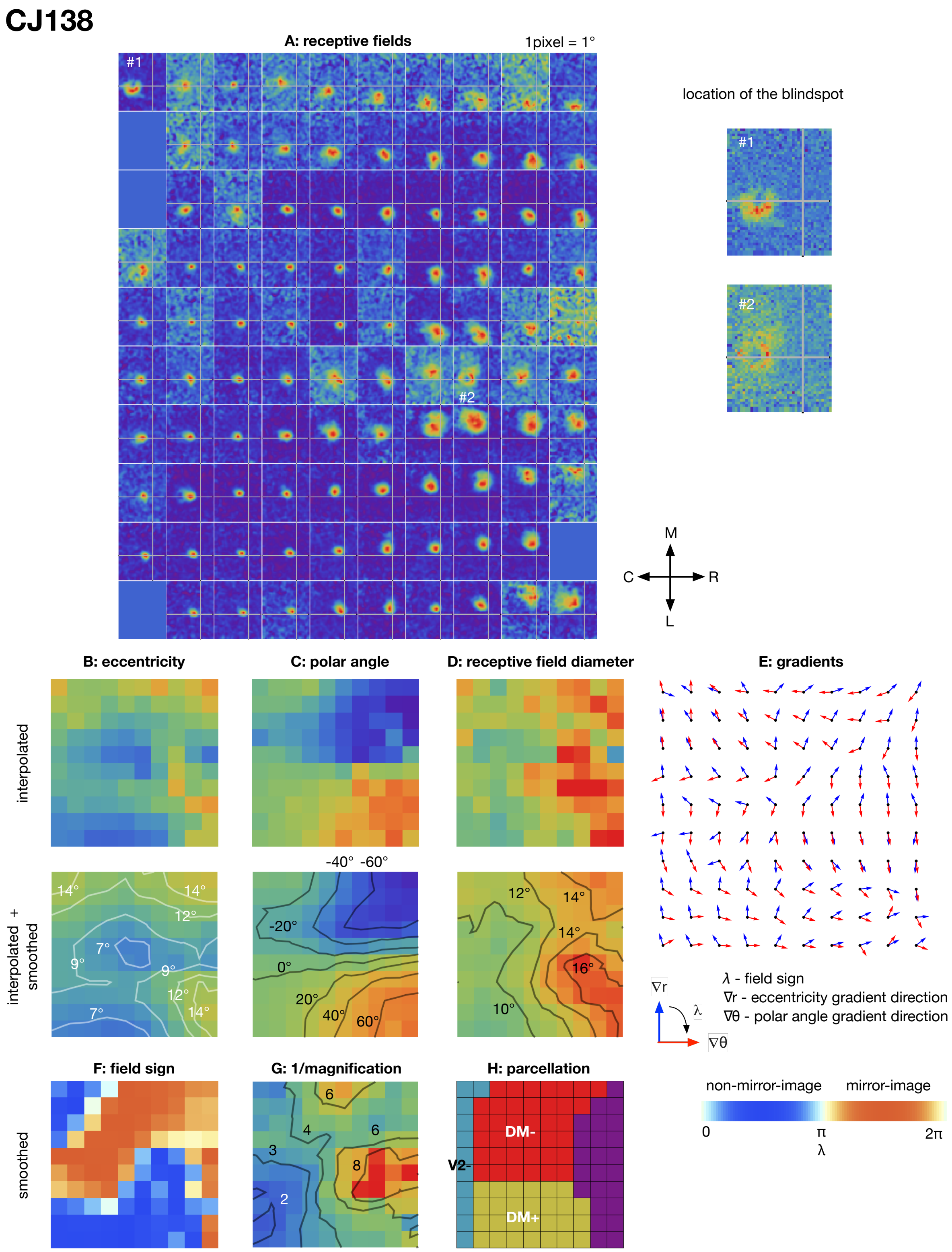


**Supplementary Figure 4**

**Summary mapping data for case CJ138**

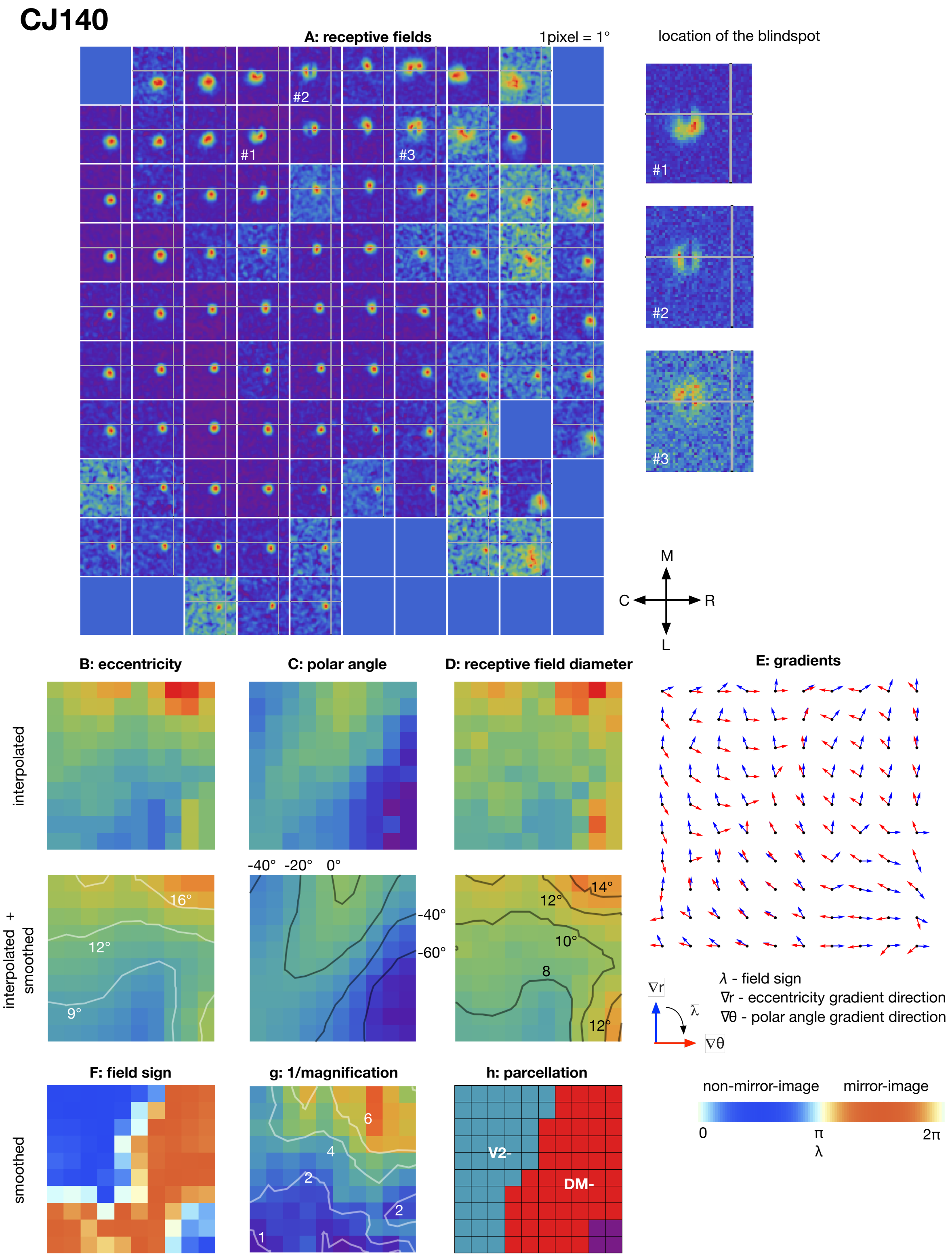


**Supplementary Figure 5**

**Summary mapping data for case CJ140**

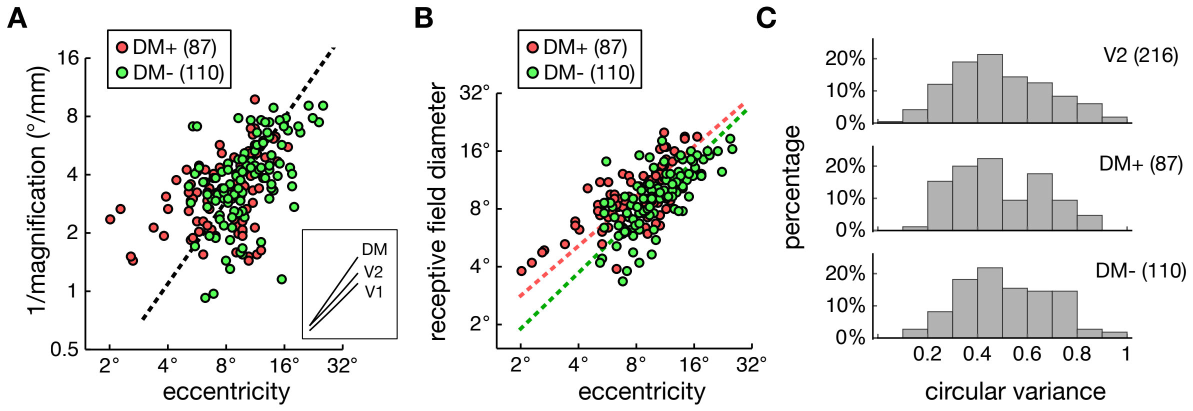


**Supplementary Figure 6**

**Comparing the response characteristics of DM+ and DM- neurons**

(**A**) The distributions of 1/M for DM+ and DM-, where M is the linear cortical magnification factor. The dashed regression line was calculated with data pooled from DM+ and DM-. Inset: The cortical magnification factor of DM compared to the marmoset V1 (Chaplin et al., 2013) and V2 (Rosa et al., 1997), in the eccentricity range of 5° to 20°.

(**B**) The distributions of receptive field diameters estimated for DM+ and DM-, as functions of the eccentricity. The dashed regression lines were calculated separately for DM+ (red) and DM- (green). (**C**) The distributions of circular variance (a measure of orientation selectivity) for V2, DM+, and DM-. Most of neurons in DM+ and DM- are orientation tuned, in a distribution similar to that of V2.

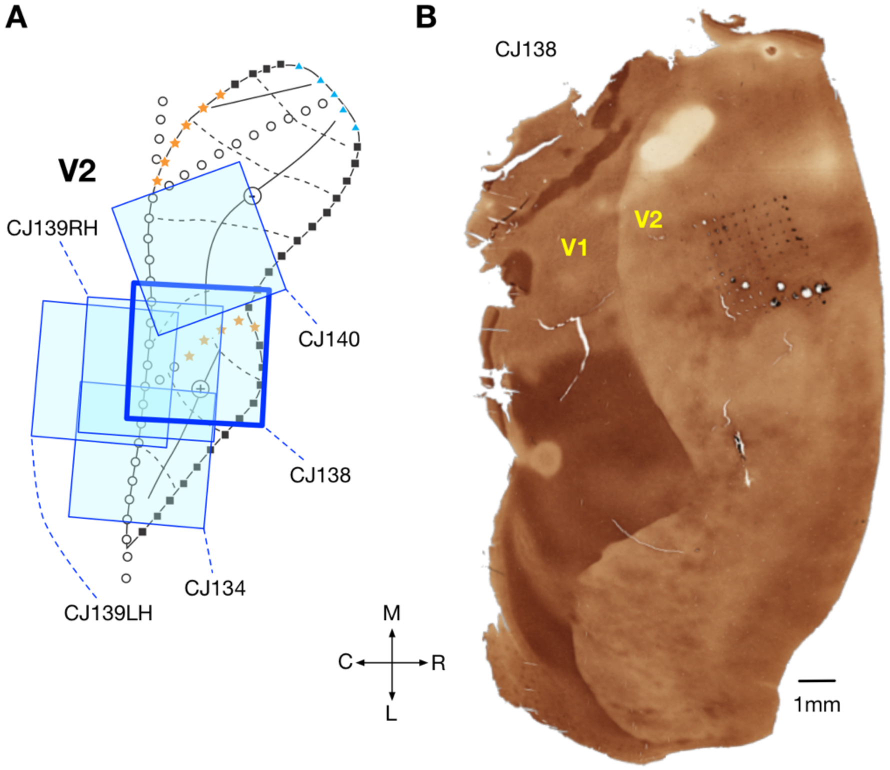


**Supplementary Figure 7**

**The locations of the implanted arrays**

**(A)** The relative locations of the implanted arrays, inferred from histology. **(B)** The flat-mounted occipital lobe (stained for cytochrome oxidase) of case CJ138. The marks left by the individual electrodes confirmed that the array was implanted immediately rostral to area V2. Abbreviations: M – medial; R – rostral; L – lateral; C – caudal.
